# Stochastic Analysis Demonstrates the Dual Role of Hfq in Chaperoning *E. coli* Sugar Shock Response

**DOI:** 10.1101/2020.06.30.178566

**Authors:** David M. Bianchi, Troy A. Brier, Anustup Poddar, Muhammad S. Azam, Carin K. Vanderpool, Taekjip Ha, Zaida Luthey-Schulten

## Abstract

Small RNAs (sRNAs) play a crucial role in the regulation of bacterial gene expression by silencing the translation of target mRNAs. SgrS is an sRNA that relieves glucose-phosphate stress, or “sugar shock” in *E. coli*. The power of single cell measurements is their ability to obtain population level statistics that illustrate cell-to-cell variation. Here, we utilize single molecule super-resolution microscopy in single *E. coli* cells coupled with stochastic modeling to analyze glucose-phosphate stress regulation by SgrS. We present a kinetic model that captures the combined effects of transcriptional regulation, gene replication and chaperone mediated RNA silencing in the SgrS regulatory network. This more complete kinetic description, simulated stochastically, recapitulates experimentally observed cellular heterogeneity and characterizes the binding of SgrS to the chaperone protein Hfq as a slow process that not only stabilizes SgrS but also may be critical in restructuring the sRNA to facilitate association with its target *ptsG* mRNA.

## Introduction

The ability of living cells to modulate their gene expression in response to changing environmental conditions is critical to their growth and continued development. Many bacteria use the phosphoenolpyruvate phosphotransferase (PTS) system to transport and phosphorylate incoming sugars to prepare them for subsequent glycolytic metabolism. The uptake of phosphosugars must be balanced with their breakdown in order to prevent metabolic stress. In *E. coli*, a stress response induced by unbalanced glucose-phosphate transport and metabolism or “sugar shock”, is referred to as glucose-phosphate stress response. A primary activity of this stress response is RNA silencing of *ptsG*, a gene coding for the glucose transport protein of the same name (also known as EIICBGlc in *E. coli*), by the small RNA (sRNA) SgrS. Small RNAs are usually non–coding RNA molecules that act by base pairing with target messengers to regulate translation or mRNA stability and have been observed across all domains of life [3]. *sgrS* is upregulated by a transcriptional activator (SgrR) when the cell is under a state of glucose-phosphate stress. SgrS regulates *ptsG* post-transcriptionally by a mechanism where SgrS binds to *ptsG* messenger RNA (mRNA) and prevents its translation to protein by blocking access of the ribosome to the mRNA [24,41]. This also enhances the co-degradation of *ptsG* mRNA and SgrS via enzymes responsible for the removal of bulk RNA such as ribonuclease E (RNase E) [20,24]. This co-degradation reduces the number of PtsG sugar transporter proteins that are produced and thus reduces the impact of glucose-phosphate stress, since fewer transport proteins are available to bring sugar into the cell.

SgrS and *ptsG* mRNA associate via complementary base pairing that occludes the ribosome binding site on the mRNA. Recently, this mechanism has been analyzed in conjunction with binding of the Sm-like chaperone protein Hfq to SgrS, which has been proposed to stabilize the sRNA, and facilitate the interaction between the sRNA and its mRNA target [18]. Hfq also promotes SgrS–dependent regulation of other targets involved in sugar shock such as *manXYZ*, and *yigL* in *E. coli*. In this study, we focus only on the primary regulatory target *ptsG* mRNA and do not consider the other targets of SgrS regulon, which are described in [6].

Previous experimental and theoretical work [19,32] has demonstrated the necessity of accounting for gene replication over the course of the cell cycle in order to capture the population variation observed in messenger RNA abundance. The additional noise emanating from transcription at multiple gene loci manifests itself in the broad mRNA copy number distributions observed in a population of cells. The aforementioned work also demonstrated that including the effect of gene regulation by transcription factors can be critical in order to appropriately describe stochastic dynamics. The effect of transcriptional regulation is apparent in the SgrS–ptsG mRNA system, where the expression of SgrS is maintained by the regulator SgrR, which activates *sgrS* and autorepresses its own expression during glucose-phosphate stress conditions [41,42].

Recently, Fei et al. [13] presented a deterministic kinetic model of the SgrS mediated regulation of *ptsG* mRNA in *E. coli*. Using single-molecule fluorescence experiments (smFISH and STORM), SgrS and *ptsG* mRNA copy numbers in cells were measured, which produced distributions of RNA at various time points after the induction of sugar stress across a population of fast-growing *E. coli*. However, it is important to note that both the *ptsG* mRNA and the SgrS regulating it are present in low copy number (a few to tens of particles) and therefore exhibit intrinsically noisy behavior in both their gene expression and regulatory behaviors. For this reason it is most appropriate to treat the regulatory network via stochastic simulation in order to quantify the variation that is observed across a population of cells, which has been demonstrated previously [11,12,34].

Here, we have developed a stochastic model, to our knowledge the first of its kind for an RNA silencing network, that captures the mRNA and sRNA distributions experimentally observed in a population of hundreds of *E. coli* cells. The stochastic model additionally incorporates the following features that extend the platform given by Fei et al [13]: (1) accounting for gene replication, (2) transcriptional gene regulation of *sgrS* by its activator SgrR and (3) explicit representation of the SgrS stabilization via the Hfq chaperone protein. This model robustly describes experimentally observed RNA distributions, closely matching regulatory dynamics from immediately after induction until a steady state is reached 20 minutes later. We also utilize this model to analyze the effects of the size of the pool of Hfq chaperone protein available to SgrS, to decouple the rate of Hfq stabilization of SgrS and its subsequent activity in enhancing association to the target, *ptsG* mRNA, and to study the effect of an *sgrS* point mutation in the SgrS-Hfq binding region on regulatory dynamics.

## Materials and Methods

### Model and Computational Methods

The previous kinetic model for SgrS regulation of *ptsG* mRNA [13] utilized simple mass-action kinetics to describe the target search process and modeled gene expression as a constitutive process, with RNA species originating from a single gene copy. Despite its simplicity, this model captures average regulatory network behavior and also gives insight into many of the parameters required for the more descriptive stochastic model that is the focus of this work. For example, since an overall binding rate for SgrS to *ptsG* mRNA was established in Fei et al. [13] we are now able to complexify the model by the addition of the chaperone protein Hfq, which allowed us to predict (by fitting to the experimental data) the size of the pool of Hfq available to stabilize SgrS and the rate at which it binds the sRNA (separate from its association to *ptsG* mRNA).

The kinetic model was implemented and solved stochastically as a well-mixed Chemical Master Equation (CME) in the Lattice Microbes (LM) simulation software suite [15,16,33,35]. The corresponding rate constants (Table 1) were adapted from the kinetic model described in Figure 1. One important feature added to the model is the explicit presence of the chaperone protein Hfq, which has been shown to both stabilize SgrS (substantially increasing its half-life) and to facilitate the association of SgrS to *ptsG* mRNA [17,36,41,43]. In order to capture the cell-to-cell heterogeneity due to the small number of particles (e.g., gene copies) involved in transcription, it is critical to account for transcriptional regulation of the genes involved in the glucose-phosphate stress response. For this reason, we include the transcriptional activation of *sgrS* by the transcription factor SgrR, which has been shown to upregulate *sgrS* expression in the presence of αMG (the unmetabolizable inducer used in place of glucose for our experiments) [41,42]. Regulation of *ptsG* by the transcriptional repressor Mlc was not included in the model since repression is relieved in the presence of glucoside sugars. With *α*MG present, Mlc is sequestered at the membrane by binding the EIIB subunit of the PtsG transporter protein complex [22,30,37], relieving repression and resulting in high levels of *ptsG* transcriptional activity [4]. Since the decay time of PtsG proteins is expected to be approximately on the order of eight hours [23], much longer than the timescale of mRNA decay, Mlc repressors are likely still sequestered by the transporters at the membrane 20 minutes post-induction and have little effect on the SgrS regulatory process. Rates for the association of the Hfq-SgrS complex to *ptsG* mRNA (*k_on_*) and the dissociation of the Hfq-SgrS-ptsG mRNA complex (*k_off_*) were obtained from [13], which did not include Hfq explicitly but provides the corresponding association and dissociation reaction rates. The value for the co-degradation rate of SgrS and *ptsG* mRNA from the Hfq-SgrS-ptsG mRNA complex by RNase E (*k_cat_*) is also obtained from Fei et al. [13] (see Section “Experimental Methods and Materials” for confirmation of *k_on_, k_off_*, and *k_cat_* values).

**Figure 1.**
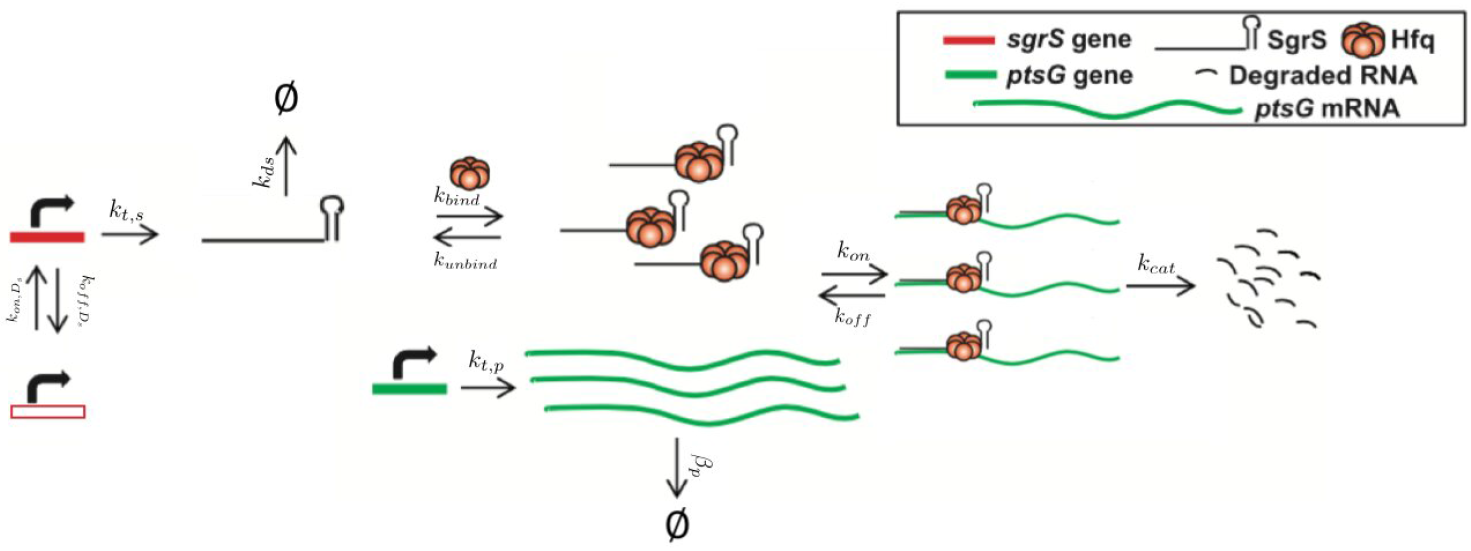
Schematic of the kinetic model as described in the text. The RNA species are transcribed from a sampled genome state with *sgrS* capable of switching between an “ON” and “OFF” state. Explicitly represented Hfq can bind and unbind with SgrS, and then the Hfq–SgrS complex binds (and potentially unbinds) with *ptsG* mRNA. All RNA degradation events are carried out by the enzyme RNase E. See Figure 4 for the kinetic equations described above.

**Table 1.**
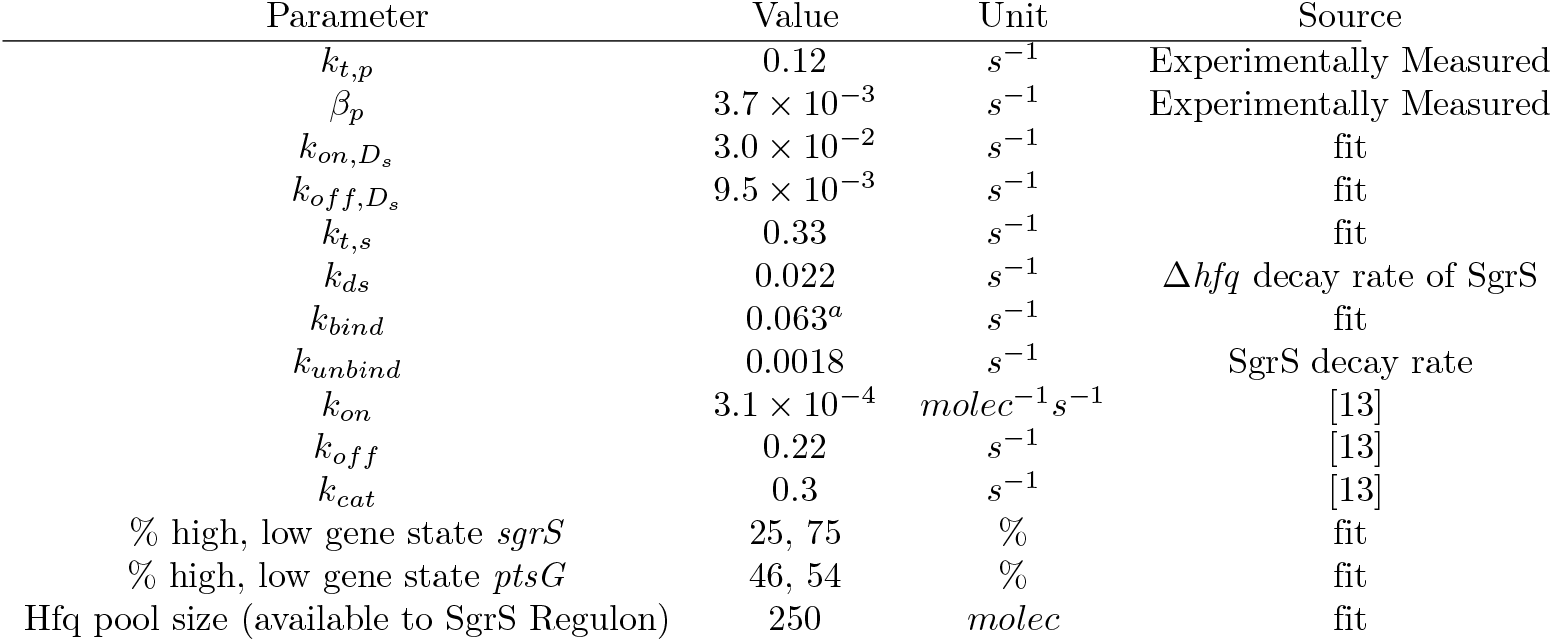
The list of parameters used for the kinetic model. The % in each gene state refers to percentage of the cellular population with the gene being in a low or high gene copy state as described in subsection. *a*) *k_bind_* is given as a Pseudo first order rate accounting for the average expected pool size of Hfq participating in SgrS stabilization and enhancement (250). When converted to the corresponding bulk second order rate with 250 Hfq present *k_bind_* agrees well with the range of Hfq binding rates measured for other sRNA reviewed in [36] and discussed further in Section “Results”

### Calculation of Gene Copy Number

Finally, and critically, in order to appropriately capture regulatory effects on gene expression of SgrS and *ptsG* mRNA, it is important to account for gene duplication, as we have previously shown [32]. As illustrated by Jones et al. [19] since the time to replicate the *E. coli* genome (approximately 40 minutes [9]) is longer than the fast-growing *E. coli* cell division time of 20 minutes (or the 35 minutes observed in our experiments), the cell has nested replication forks that are already replicating the genomes of daughter and granddaughter cells prior to cell division. In particular, genes close to the origin of replication are likely to have multiple copies present over much of the cell cycle. This phenomenon has been shown previously for genes near the origin in *E. coli* by both isotopic labeling of nucleotides and imaging of fluorescent chromosome markers [9,48]. Due to the position of *sgrS* (only 6° away along the circular chromosome) very near to the origin of replication, it is likely that multiple gene copies are accessible for transcription over the course of the cell cycle. About half-way between the origin and terminus of replication (at approximately 90°) *ptsG* is also likely to have multiple gene copies present at some point over the course of the cell cycle, although at lower copy number than *sgrS*. Figure 2 depicts the two genes and their location along the circular *E. coli* genome. The experimentally measured cells were unsynchronized and should have multiple replication forks present over the course of the 20 minutes post-induction, our measurement window. To account for gene duplication effects in a population of unsynchronized cells, we sample the percentage of the cellular population in either a low or high gene state, which corresponds to the expected distribution of the number of genes present over the course of the cell cycle after induction. In this way, we effectively flip a coin to decide whether a simulation replicate corresponding to an individual experimentally imaged *E. coli* cell has 2 copies (low gene state) or 4 copies (high gene state) of *sgrS* and similarly 1 or 2 copies of *ptsG*. This allows us to account for the effect of gene duplication in generating mRNA noise over the heterogeneous population of hundreds of *E. coli* cells that were observed experimentally. We assume that all gene copies are transcribed independently from one another and at the same rate, a notion that [46] has recently examined in *E. coli* under various growth conditions.

**Figure 2.**
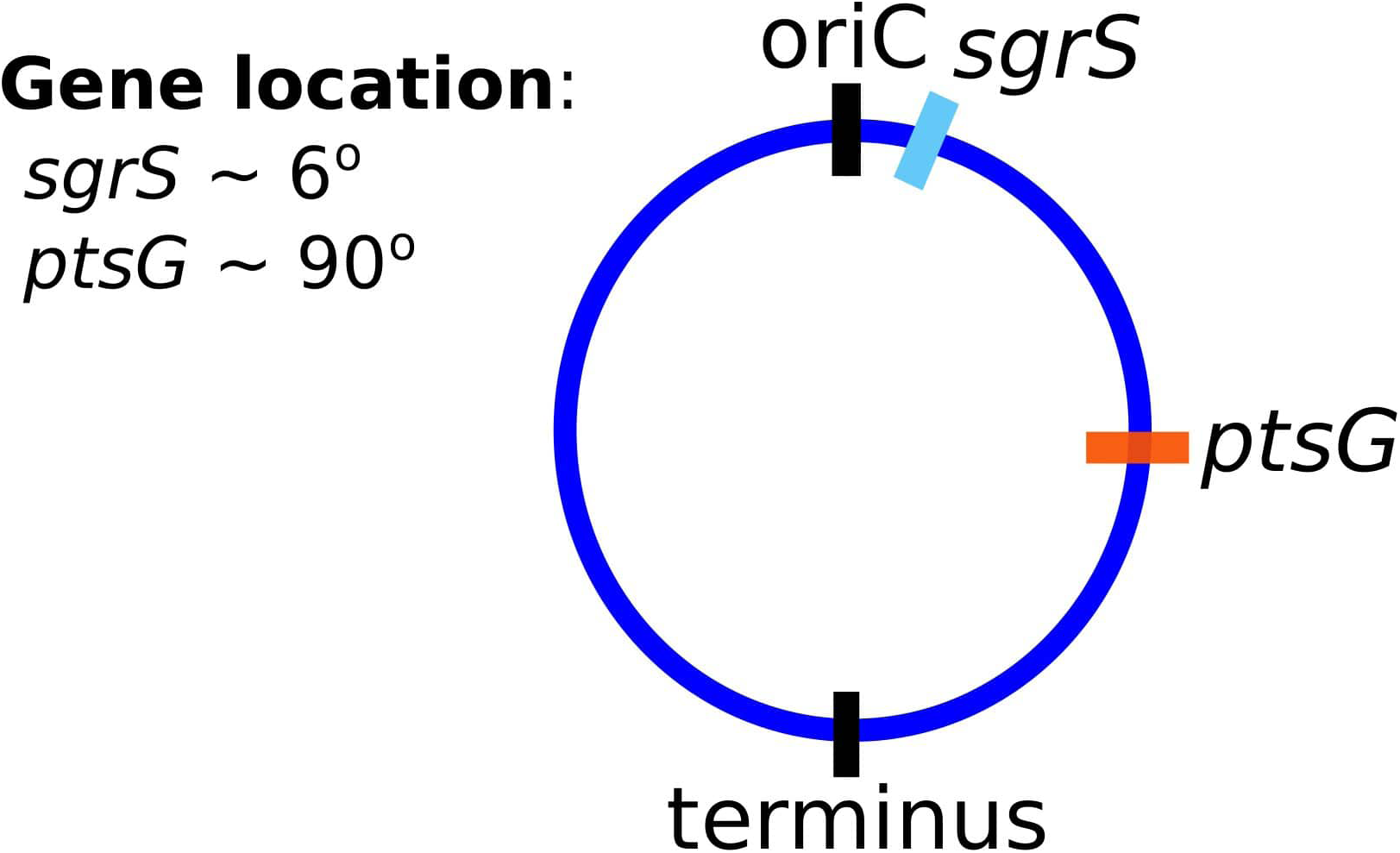
The gene location for SgrS and *ptsG* mRNA relative to the origin of replication (*oriC*) are shown on the circular genome of the *E. coli* cells used for this study. As it is closer to the origin of replication *sgrS* (cyan) is likely to be present in higher gene copy number than *ptsG* (orange), which is farther away from the *oriC*.

Under similar growth conditions to ours (MOPS glucose-based medium with a doubling time of 35 minutes, (see Section “Experimental Methods and Materials”)), the data from [46] suggest that transcription does appear to be independent and uncorrelated between copies of the same gene. Figure 3 illustrates the reasoning for the specific choices of high and low state gene copy numbers for *ptsG* and *sgrS* in an *E. coli* cell growing faster than the expected time necessary for replication (approximately 40 minutes, compared to an experimentally observed generation time of approximately 35 minutes) [9,48]. Stochastic simulations were performed by sampling the CME for the model given in Figure 1 with the widely used Gillespie Direct Method of the Stochastic Simulation Algorithm (SSA), which is implemented in the publicly available Lattice Microbes (LM) software suite (version 2.3 was used) and its python interface pyLM [15,16,33,35]. We ran 2000 replicate simulations for 25 minutes after *α*MG induction of glucose-phosphate stress in order to match the corresponding 20-minute smFISH-STORM experiments. Initial conditions for basal SgrS (1-3 copies) and *ptsG* mRNA (30-40 copies) copy number were sampled from the experimentally measured distributions and rounded to the nearest integer particle number (a necessity for stochastic representation). Simulations were computed on a local cluster containing AMD Opteron Interlagos cores.

**Figure 3.**
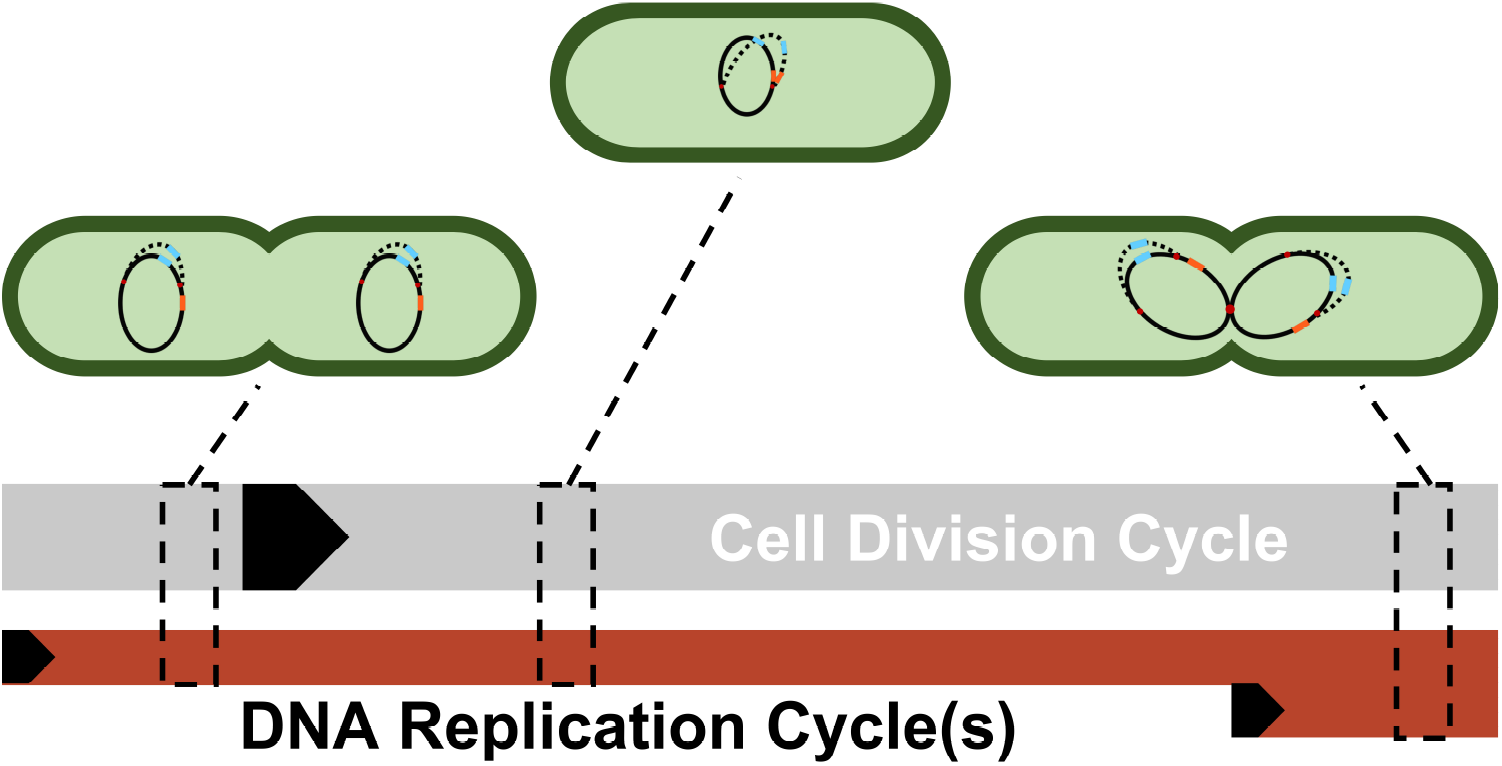
A simplified depiction of possible cellular states throughout a single DNA replication cycle. Each cell shows a snapshot of the gene state of a cell given its progression through the DNA replication and cell division cycle. Due to the difference in lengths of the cell division cycle (~35 mins) and DNA replication cycle (~40 mins), DNA replication and cell division are not completely in sync. Multiple replication forks (red dots) can form on the genome in order to ensure DNA is duplicated properly in these fast-growing cells. As a result, genes closer to the origin such as *sgrS* (blue) are duplicated in the same timeframe that replication is initiated (resulting in 2 or 4 gene copies), while genes closer to the terminus such as *ptsG* (orange) are replicated in the C period, the period when a majority of DNA is duplicated (resulting in 1 or 2 gene copies). The black arrows denote the start of a cycle.

### SgrS Regulatory Network Kinetic Model

The kinetic model describing the reactions characterizing the *E. coli* glucose-phosphate response network by the small RNA SgrS is given in Figure 4. Simulation files are available in Jupyter Notebook format to be simulated via the Lattice Microbes (LM) Software Package at **Add link to jupyter notebook**.

**Figure 4.**
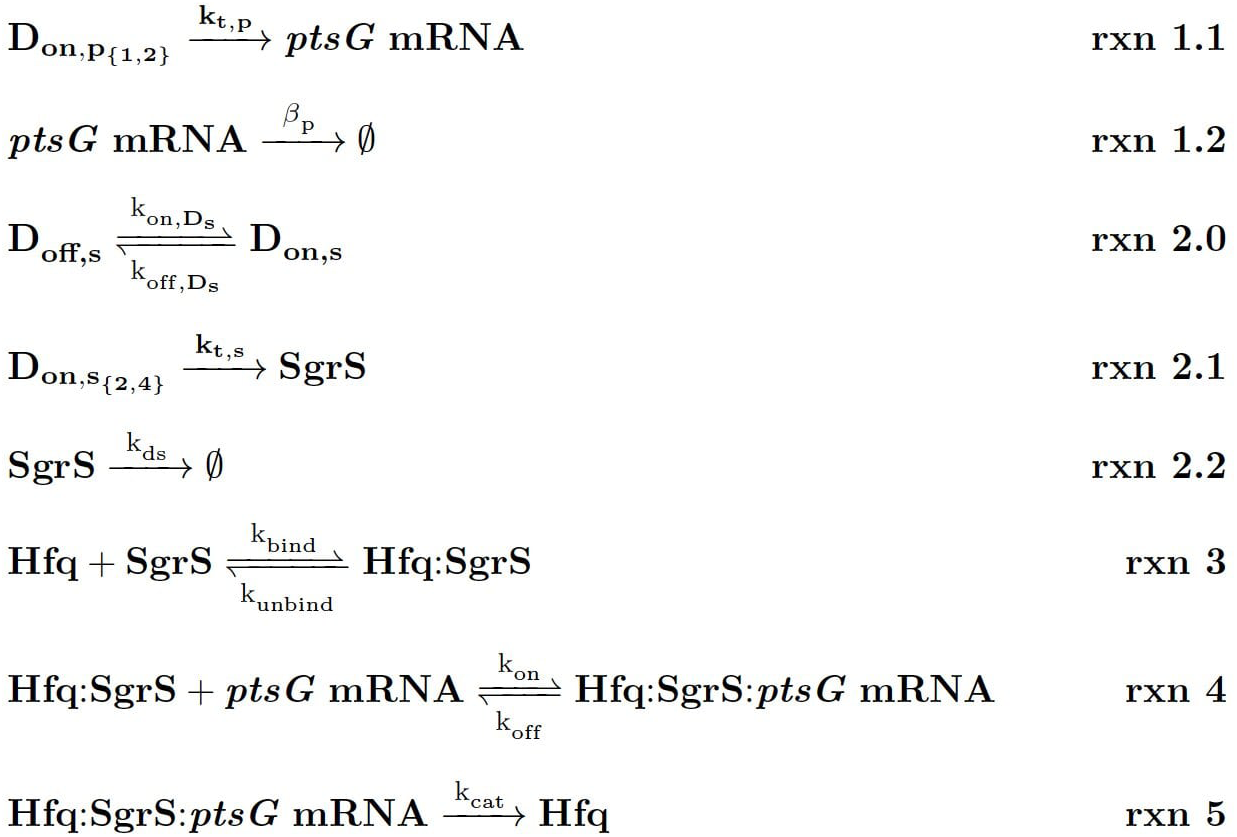
Kinetic Equations of the SgrS regulatory network. *D*_*on,p*_1,2__ refers to the gene (or DNA) for *ptsG* in 1 (low state) or 2 (high state) copies and *D*_*on,s*_2,4__ corresponds to the gene for *sgrS* in 2 (low state) or 4 (high state) copies. *D*_*on,s*_ corresponds to *sgrS* when it is in the “ON” state due to activated or solute bound transcriptional activator SgrR being bound [42]. *k_ds_* corresponds to the experimentally measured degradation rate of SgrS when cellular Hfq is not present and *k_unbind_* corresponds to the experimentally measured degradation of SgrS when Hfq was present.

### Experimental Methods and Materials

Wild type *E. coli* cells (DJ480) were grown overnight at 37 °C, 250 rpm in LB Broth. The SgrS U224G mutant was grown in LB Broth with 50 *μ*g/ml spectinomycin (Spec) (Sigma-Aldrich). The next day, overnight cultures were diluted 100-fold into MOPS EZ rich defined medium with 0.2% glucose and the cells were grown until *OD*_600_ reached 0.15–0.25. *α*-methyl D-glucopyranoside (*α*MG) (Sigma Aldrich) was then added to provoke glucose-phosphate stress and induce SgrS expression response. Specific volumes of liquid were removed from the culture at 0, 2, 4, 6, 8, 10, 15, and 20 minutes after induction and mixed with formaldehyde (Fisher Scientific) to a final concentration of 4% for cell fixation prior to single molecule experiments. Following fixation, the cells were incubated and washed, before being permeabilized with 70% ethanol, to allow for fluorescence *in situ* hybridization (FISH). Stellaris Probe Designer was used to design the smFISH oligonucleotide probes that were ordered from Biosearch Technologies (https://www.biosearchtech.com/). Each sRNA was labeled with 9 Alexa Fluor 647 probes while each *ptsG* mRNA was labeled with 28 CF 568 probes. The labeled RNA molecules were then imaged via the super-resolution technique STORM (Stochastic Optical Reconstruction Microscopy). A density-based clustering analysis algorithm (DBSCAN) [10] was utilized to calculate RNA copy numbers. The algorithm used was the same as previously published [13], but the Nps and Eps values were updated for the SgrS and *ptsG* mRNA images, since CF 568 was used instead of Alexa Fluor 568 and a 405 nm laser was used to reactivate the dyes. The SgrS (9 probes labeled with AlexaFluor 647) images were clustered using *Nps* = 3 and *Eps* = 15 and the *ptsG* mRNA (28 probes labeled with CF 568) images were clustered using *Nps* = 10 and *Eps* = 25 and these numbers were empirically chosen. A MATLAB code was used for cluster analysis. The raw data was acquired using the Python-based acquisition software and it was analyzed using a data analysis algorithm which was based on work previously published by [2]. The peak identification and fitting were performed using the method described previously [13]. The z-stabilization was done by the CRISP system and the horizontal drift was calculated using Fast Fourier Transformation (FFT) on the reconstructed images of subsets of the super-resolution image, comparing the center of the transformed images and corrected using linear interpolation. The *ptsG* mRNA degradation rates were calculated via a rifampicin-chase experiment. The wild type (DJ480) *E. coli* cells and Δ*hfq* mutant strain SA1816 [DJ480, *laclg, tetR, spec*, Δ*hfq::kan*] cells were grown in LB Broth with the respective antibiotics at 37 °C, 250 rpm overnight. They were used to calculate the RNA degradation rates. The Δ*hfq::kan* allele was moved to create strain SA1816 constructed by P1 transduction [27]. When the *OD*_600_ reached 0.15–0.25, rifampicin (Sigma-Aldrich) was added to cultures to a final concentration of 500 *μ*g/ml. The cells were labeled by smFISH probes and analyzed by the same process described above, taking the time of rifampicin addition or *α*MG removal as the 0 time point. Aliquots were taken after 0, 2, 4, 6, 8, 10, 15, and 20 minutes (0, 2, 4, 6, and 8 minutes for ΔHfq strains). For the purpose of background subtraction, ΔSgrS and Δ*ptsG* mRNA strains were grown, labeled with probes and imaged in the same manner to be used for the measurement of the background signal due to the non-specific binding of Alexa Fluor 647 and CF 568. The natural logs of the RNA copy numbers were plotted against time and the slope of the linear fitting was used to calculate the RNA lifetime and then the degradation rates. SgrS degradation rates were obtained from [13], where they were measured by stopping the transcription of *sgrS* by removing *α*MG from the media and then were imaged and analyzed to calculate the degradation rates in the same manner as was described for *ptsG* mRNA. The values for *k_cat_, k_on_*, and *k_off_* for WT cells were confirmed to be within the errors reported for the values given in [13] by fitting to the experimentally measured RNA counts with the simplified model given in that work. The transcription rate of *ptsG* was determined using *k_t.p_* = *β_p_* × [*p*]_0_, (as described in [13]), where [*p*]_0_ was the average initial level of *ptsG* mRNA before stress induction. The transcription rate obtained was unchanged between the wild-type and the U224G mutant cells.

## Results

Figure 5 demonstrates the ability of our newly developed kinetic model to capture the average cellular copy number of SgrS and *ptsG* mRNA over the course of the 20 minute period post-induction. The overlap of the interquartile range (IQR) of both the experimental and simulated cellular populations demonstrates the agreement over a variety of cells (at different gene states (*i.e* high/low copy number), and RNA expression levels).

**Figure 5.**
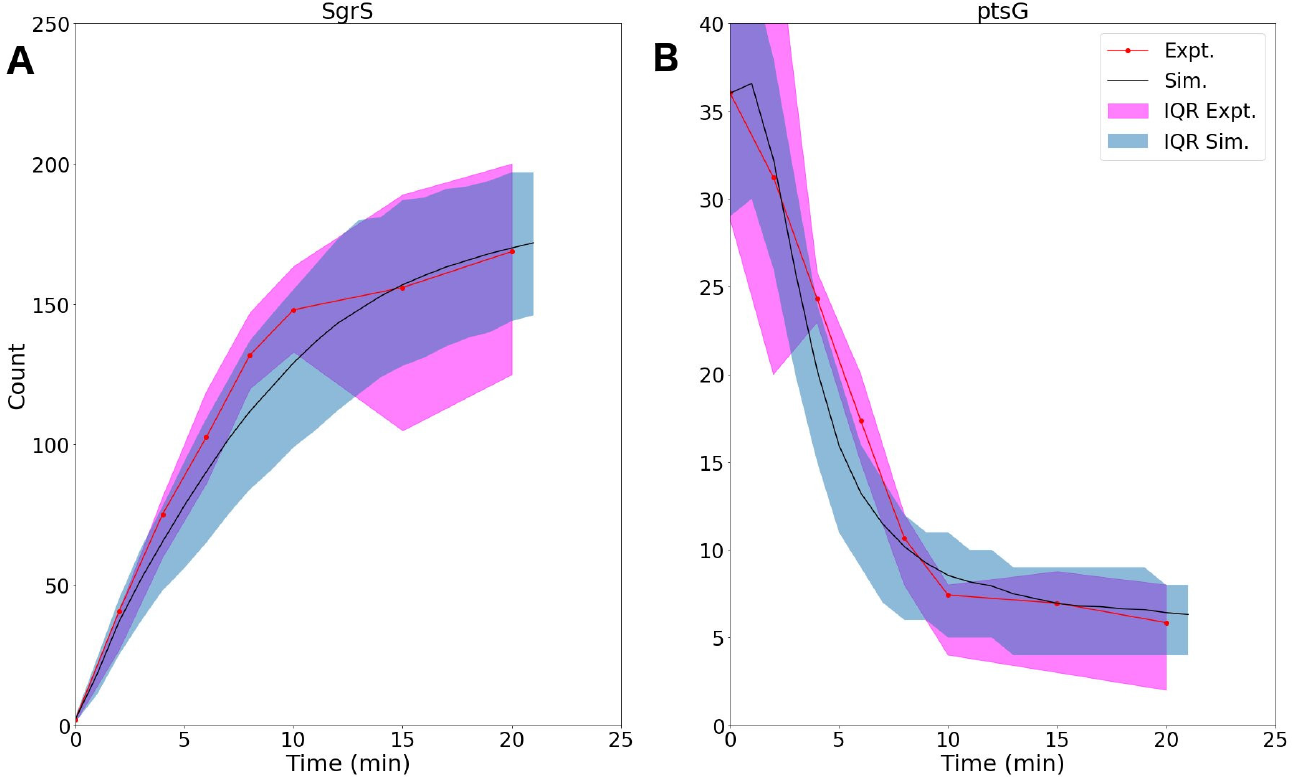
Average time trace and interquartile range (IQR) of **(A)** labeled SgrS and **(B)** *ptsG* mRNA from both 85–169 cells from smFISH experiments (red) and 2000 replicates from kinetic model simulations (blue). The kinetic model shows strong agreement, especially at long times (10-20 minutes) after induction and captures overall response behavior. An available pool of 250 Hfq and the kinetic parameters given in Table 1 were utilized. Results considering both lower and higher available Hfq pools are discussed in **Supplemental File 1–Figure 9**.

The ability of our improved kinetic model to capture population-level statistics of single cell copy number distributions of SgrS and *ptsG* mRNA is demonstrated in Figure 6. Kernel Density Estimates (KDE), which are used to estimate the probability densities of distributions of approximately 100–200 experimentally measured cells and 2000 simulated cells are displayed, along with dashed vertical lines giving the average RNA copy numbers observed. KDEs were utilized to provide a reasonable comparison to the experimental values despite the fact that there were a relatively low number of cells measured at each time point (approximately 100–200) compared to the number of replicates required for appropriate stochastic simulation (2000) (Histograms of experimental RNA counts measured before KDE imposition are given in **Supplemental File 1–Figure 15**, Figure 14). The distributions obtained from both experiment and the kinetic model show strong agreement (especially in the case of *ptsG* mRNA), which can be seen quantitatively by the starred line showing the Kulback–Leibler Divergence (KL Divergence) in Figure 7. The KL Divergence (Equation 2), which was minimized to fit to experimental RNA distributions over all time points, is a statistical measure used to characterize the difference between a probability distribution (the KDE of simulated cells) and a reference distribution (the KDE of experimentally measured cells).

**Figure 6.**
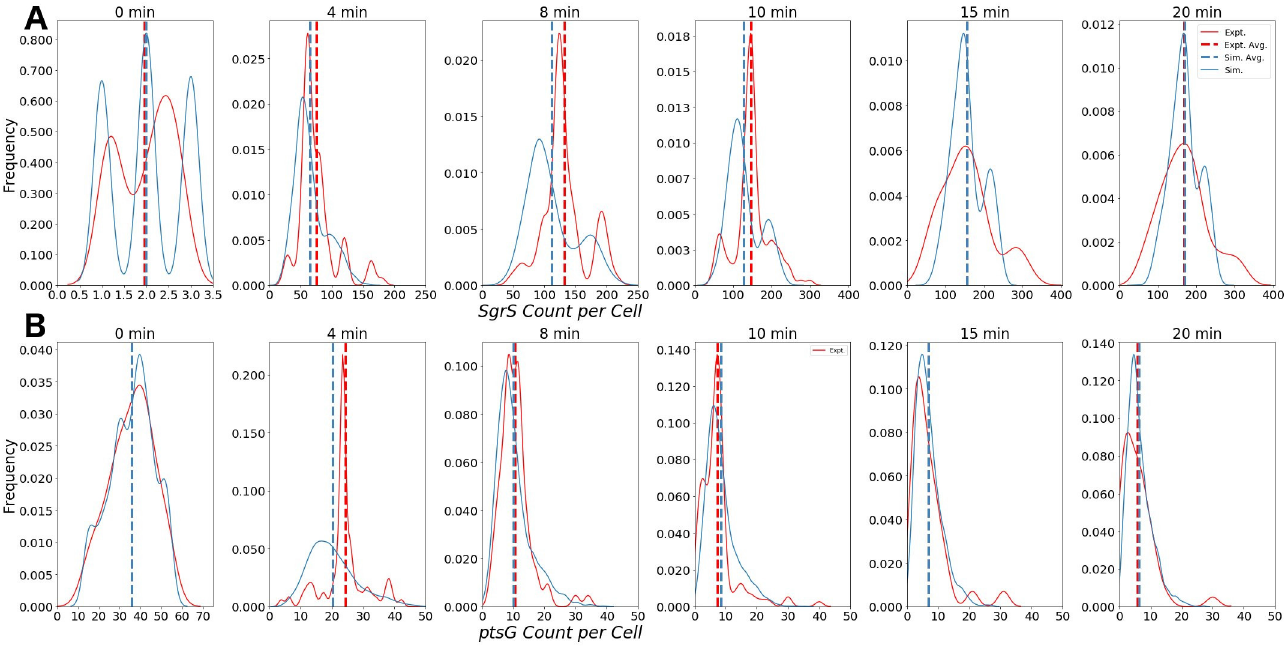
Distributions of **(A)** Wild-Type SgrS (top) and **(B)** *ptsG* mRNA (bottom) at various time points from 0 to 20 minutes post-induction. Data from smFISH-STORM experiments (red, 100–200 cells per time point) and stochastic simulations (blue, 2000 cells per time point) are shown as kernel density estimates. The effect of number of cell replicates is studied further in **Supplemental File 1–Figure 11**. Average copy number at each time point is are displayed with dashed vertical lines.

**Figure 7.**
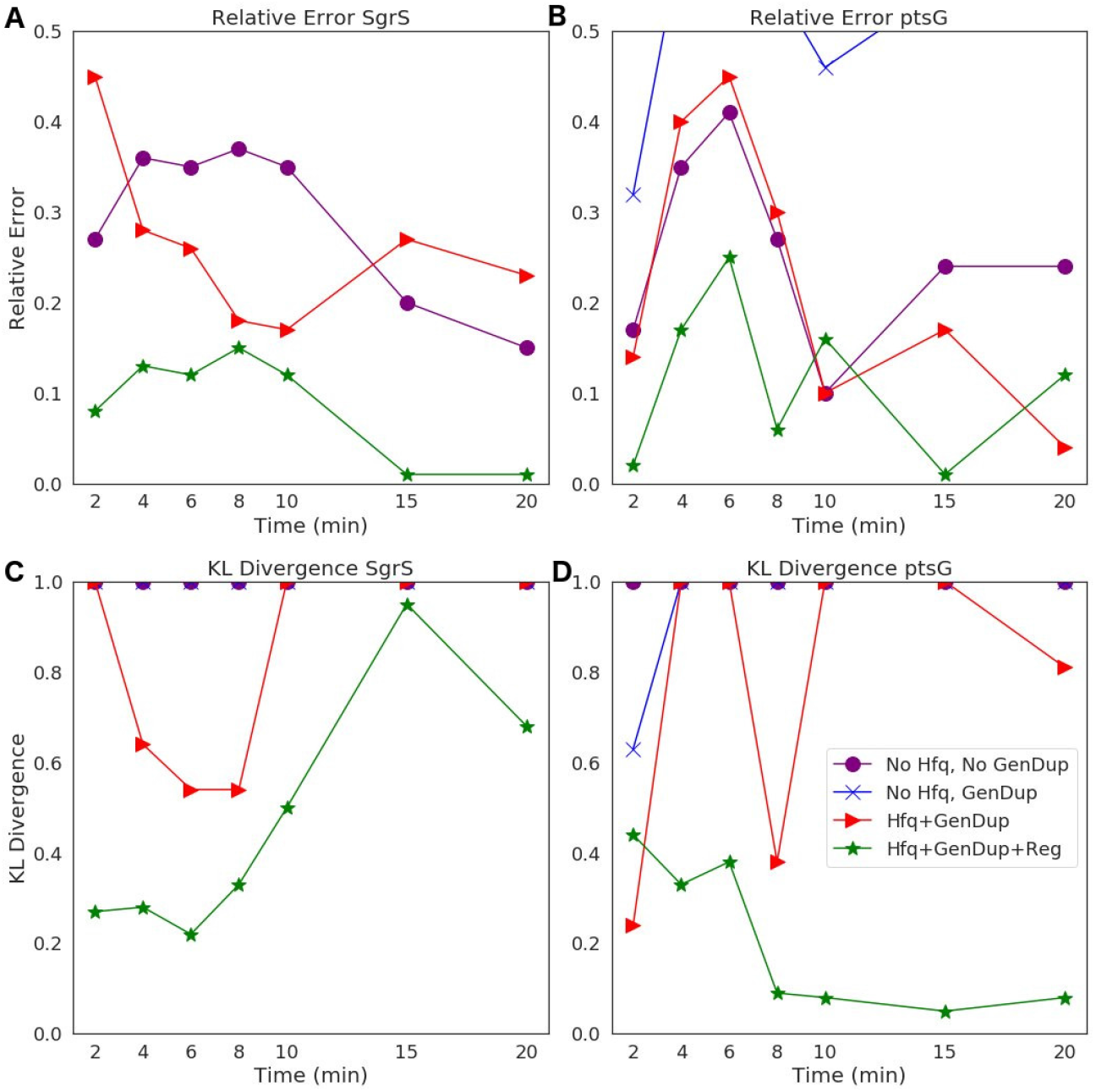
Statistical analysis of the agreement of **(A),(B)** *SgrS* sRNA and **(C),(D)** *ptsG* mRNA copy number between experiment and theory on both **(A),(C)** an average (Relative Error) and **(B),(D)** distribution (Kulback–Leibler: KL Divergence) level. KL Divergence values for the model with no Hfq stabilization nor Gene Duplication are not shown as the values obtained are at 1.0, corresponding to significant disagreement in that model variant and experiment. GeneDup refers to a model with Gene Duplication for both SgrS and *ptsG* implemented and Reg refers to a model with transcriptional regulation of SgrS by SgrR in place. The green line (with star markers) indicates the full kinetic model used for this study, which provides the best fit to both average and population level data for both SgrS and *ptsG* mRNA.

The parameters obtained from the fitting process give some insight into the role of stabilization by Hfq in the SgrS-*ptsG* mRNA target search process and the role of transcriptional regulation by SgrR in the regulatory network. The pseudo first order rate of Hfq binding to SgrS (*k_bind_*) is 0.063 s^−1^, while the degradation rate of SgrS (*k_ds_*), obtained from Δ*hfq* strain experiments (described in Section “Experimental Methods and Materials”), is 0.022 s^−1^. The available Hfq pool size of 250 predicted by fitting to the kinetic model seems reasonable in that average proteomics values have been found to be on the order 1500 [36, 40] and unique sRNAs have been shown to be bound to 10 to 1000 copies of Hfq in *E. coli* [26] (**Supplemental File 1–Section 1**). Additionally, the aforementioned SgrS-Hfq binding rate *k_bind_* corresponds well to experimentally measured *in vitro* values for sRNA-Hfq binding for sRNA of its approximate size [14,17,36]. If the pseudo first order rate for *k_bind_* reported in Table 1 is converted to a bulk second order rate by incorporating the Hfq concentration at the predicted available pool size of 250, we obtain a binding rate of 1.5 × 10^5^ *M*^−1^ *s*^−1^. This value agrees better with the reported value of approximately [36] 10^6^ *M*^−1^ *s*^−1^ for long RNAs binding to Hfq [14,21] than 10^8^ *M*^−1^ *s*^−1^ reported for short, unstructured RNAs binding to Hfq [17]). Since SgrS is a relatively long sRNA (sRNA have typically been found to be between 37–300 nt [44]) with a length of 227 nucleotides, the slow sRNA-Hfq binding rate obtained by fitting seems appropriate. This type of slow sRNA association process has been suggested to be characterized by RNA restructuring (by which Hfq remodels sRNA regions in order to make its secondary structure more accessible for target mRNA base pairing) [1,8,38,39], which has been proposed to occur for SgrS [24]. *k_bind_* is also much greater than the Hfq-SgrS unbinding rate (*k_unbind_*) of 0.0018 *s*^−1^ which was obtained from fitting to the degradation rate of SgrS in a cell where Hfq was expressed (distinct from the Δ*hfq* rate) by assuming that Hfq-SgrS unbinding is the rate-limiting step in the degradation of free SgrS represented in Figure 4 Reaction 2.2. These results seem reasonable in that SgrS should associate with Hfq at a rate comparable to its degradation as well as that SgrS-Hfq binding should happen at a significantly higher rate than their dissociation for sRNA chaperone stabilization by Hfq to be effective.

The kinetic values for transcriptional regulation by the activator SgrR also seem reasonable with a *k_on,Ds_* of 3.0 × 10^−2^ *s*^−1^ and a *k_off,Ds_* of 9.5 × 10^−3^ *s*^−1^. The gene switching parameters correspond to *sgrS* activation via SgrR binding occurring approximately 30 seconds after initiation of induction, with all *sgrS* genes assumed to start in the “OFF” state (the effect of starting genes in the “OFF” versus the “ON” state is analyzed in **Supplemental File 1–Figure 10**). This seems reasonable since SgrS sRNA moves from a basal level of a few copies to greater than 40 copies on average in two minutes time (Figure 5). The fact that *k_on,Ds_* is 3 times greater than *k_off,Ds_* means that activation happens more frequently than deactivation from unbinding of SgrR. This relative behavior is somewhat expected as sugar shock has been induced and SgrR is believed to be transformed to its active conformation as a transcriptional factor for *sgrS* by binding to a small molecule at its C-terminus [41,42]. While the available evidence suggests that the activity of SgrR due to solute binding rather than *sgrR* expression affects activation of *sgrS*, it has been demonstrated that SgrR is negatively autoregulated [42] which may lead to a ceiling on the level of *sgrS* activation that can occur even after glucose-phosphate stress is fully induced. Thus, we incorporate constant rates of *k_on,Ds_* and *k_off,Ds_* for *sgrS* activation in our model, instead of a time variant rate constant for either parameter.

### Comparison of Goodness of Fit Based on Model Complexity

To illustrate the improvement of the kinetic model to describe cellular populations, we compare simulation results sequentially as each level of complexity (*i.e*., transcriptional regulation by SgrR, gene replication, and stabilization by the chaperone protein Hfq) is added to the original reduced model presented in [13]. Figure 7 demonstrates the improvement in descriptiveness at both an average and population level with progression to a more fine-grained kinetic model. The relative error (Equation 1) of the average copy number of SgrS and *ptsG* mRNA gives the capability of the model to reproduce experiments on an average level, while the Kulback-Leibler Divergence (KL Divergence) (Equation 2) shows the agreement between the experimentally observed and simulation distributions of RNA copy numbers at a series of times from 0 to 20 minutes post induction.

The Relative Error used to illustrate the agreement between the experimentally measured average RNA copy number and the theoretical value is given by:

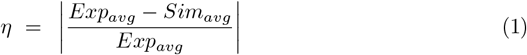

The KL Divergence used to compare agreement between experimental and simulated distributions is given by:

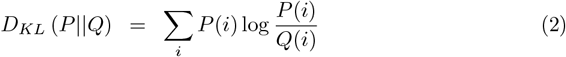

where P(i) is the continuous probability distribution given by the Gaussian KDE of the experimental copy number distribution of RNA (SgrS or *ptsG* mRNA) and Q(i) is the analogous simulated RNA copy number distribution.

It is clear that the decrease in the KL Divergence (Figure 7C,D), describing the ability of the kinetic model to accurately describe cell-to-cell variation, is most substantial in the final model presented in this work (star markers). Accounting for transcriptional regulation by SgrR, ongoing gene replication, and the stabilizing effect of Hfq allows for a more faithful description of the noise observed in a cellular population in the process of sugar shock response.

### Characterizing the Effects of SgrS Point Mutation on Association to Hfq and ptsG *mRNA*

The stochastic model we have presented can also be utilized to characterize the effects of *sgrS* point mutations on the regulatory network as a whole. The polyU tail region of *sgrS* comprising the final 8 residues of the 5’ end (all of which are uridine in the sRNA) has previously been shown to be an important site for Hfq recruitment [31]. When the polyU tail is truncated or similarly disrupted, there is a noticeable decrease in SgrS regulatory efficiency. With this in mind, we used the previously defined kinetic model (See Figure 4) to characterize the effect of a point mutation resulting in a U to G change in SgrS at position 224 (in the polyU tail region, hereafter referred to as U224G) of the sRNA on regulatory kinetics. This point mutation is well downstream of the seed region (nucleotides 168–187) where SgrS-*ptsG* mRNA base pairing occurs [7,24] so it should not directly interfere with sRNA-mRNA interactions. It is also important to consider the possible structural effects arising from polyU tail mutation. Through *in silico* folding with the RNA structure prediction tool mFold [49], we confirmed that the stability of the U224G with a ΔG of – 17.60 *kcal/mol* is unchanged from the predicted wild-type value of – 17.60 *kcal/mol*, and also indicated that sRNA structure is conserved **Supplemental File 1–Figure 13**) and the measured wild-type ΔHfq degradation rate (see Section “Experimental Methods and Materials”) is appropriate for use in fitting the U224G mutant data (as a rate for Figure 1, **rxn 2.2**).

We then fit to the experimentally measured SgrS and *ptsG* mRNA distributions using the previously determined kinetic model. A robust fit describing both average behavior as well as population level variation (Figure 8, **Supplemental File 1–Figure 12**) was achieved primarily by modulating the rates of SgrS to Hfq binding and unbinding and the *ptsG* mRNA annealing rates *k_on_* and *k_off_* (which were also free parameters in this treatment) to a much lesser extent, which further demonstrates the role of the polyU tail in Hfq chaperone recruitment.

**Figure 8.**
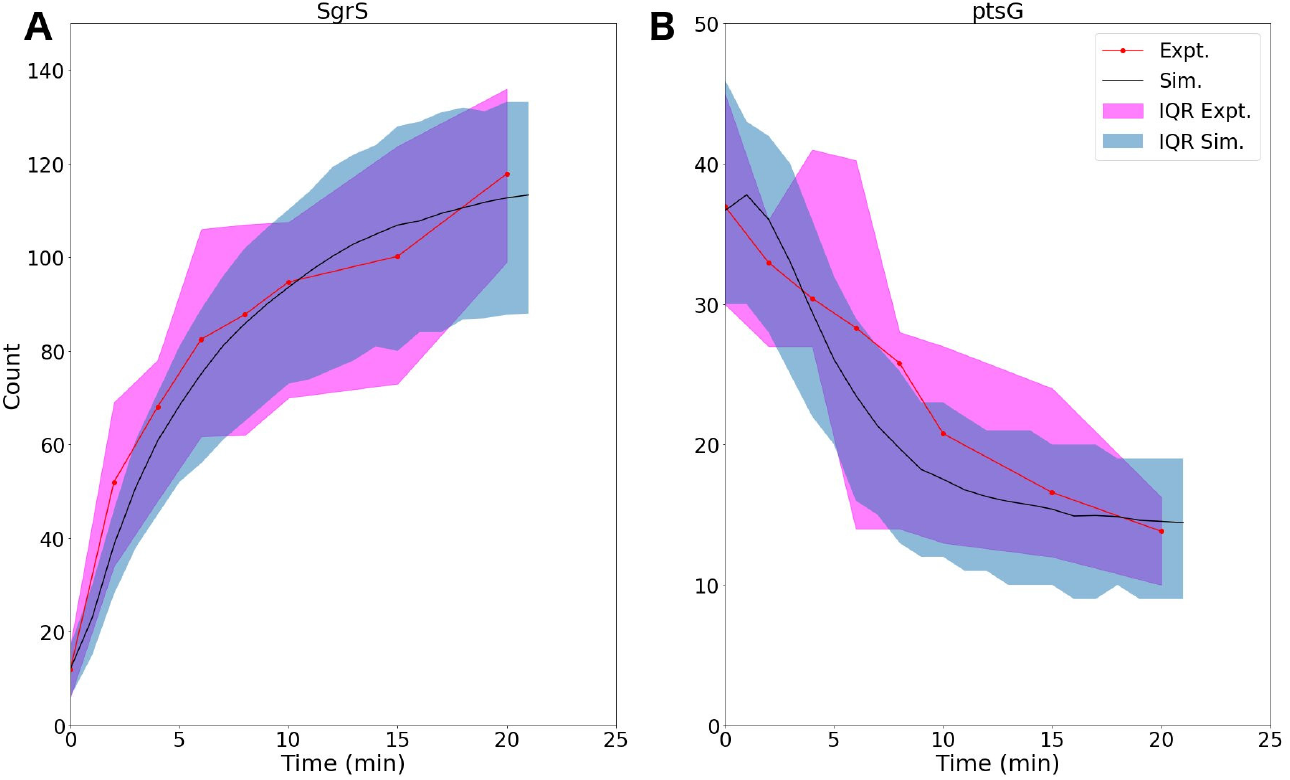
For U224G mutant cells, average time trace and interquartile range (IQR) of **(A)** labeled SgrS and **(B)** *ptsG* mRNA from both 85-169 cells from smFISH experiments (red) and 2000 replicates from kinetic model simulations (blue). The kinetic model shows strong agreement, especially at long times (10-20 minutes) after induction and captures overall response behavior. An available pool of 250 Hfq and the kinetic parameters given in Table 1 were utilized, other than changes to SgrS-HFq binding and unbinding rates and *ptsG* mRNA annealing and dissociation rates given in Table 2.

The changes in the kinetic parameters of the model used to fit mutant U224G relative to the wild-type cells (WT) illustrate that the effects of this mutation on SgrS-Hfq association are much larger, relative to the subsequent annealing of SgrS to its target *ptsG* mRNA (Table 2) (Futher discussion in **Supplemental File 1–Section 4**).

**Table 2.** The list of kinetic parameters for SgrS-Hfq association (*k_bind_* and *k_unbind_*) and annealing with *ptsG* mRNA (*k_on_* and *k_off_*) for wild-type (WT) cells as well as SgrS mutant U224G (Reactions in Figure 4). The substantial decrease in the values of *k_bind_* and *k_unbind_* demonstrate the disruption of Hfq binding that accompanies the mutation in the polyU tail, which has been observed previously [31]. The smaller relative changes in the *ptsG* mRNA annealing rates may be due to disruption of RNA restructuring [1,8,38,39] of SgrS by Hfq that hampers association to the mRNA target.

| Parameter | Mutant | Value | % Difference from WT |
| --- | --- | --- | --- |
| $k_{bind}$ | U224G | $0.033 \text{ s}^{-1}$ | -48% |
| | WT | $0.063 \text{ s}^{-1}$ | |
| $k_{unbind}$ | U224G | $0.003 \text{ s}^{-1}$ | +66% |
| | WT | $0.0018 \text{ s}^{-1}$ | |
| $k_{on}$ | U224G | $2.1 \times 10^{-4} \text{ molec}^{-1} \text{ s}^{-1}$ | -32% |
| | WT | $3.1 \times 10^{-4} \text{ molec}^{-1} \text{ s}^{-1}$ | |
| $k_{off}$ | U224G | $0.27 \text{ s}^{-1}$ | +22% |
| | WT | $0.22 \text{ s}^{-1}$ | |

The 48% decrease in the SgrS-Hfq binding rate *k_bind_* and 66% increase in the unbinding rate of the sRNA and chaperone complex *k_unbind_* highlight the effects of polyU tail disruption, and support previous conclusions that this is an important site for Hfq stabilization of SgrS [31], and the regulatory efficiency of the network as a whole. The smaller relative changes in the SgrS-ptsG mRNA annealing rates *k_on_* and *k_off_* by 32% and 22% respectively may be due to altered interactions with Hfq that impair Hfq–dependent annealing of SgrS and *ptsG* mRNA (**Supplemental File 1–Section 4**). In light of the previously discussed slow SgrS–Hfq association process, it is reasonable that RNA restructuring of Hfq may be disrupted by mutation U224G, thus leading to slower and weaker annealing to *ptsG* mRNA. One possible explanation for the disturbance of regulation in mutant U224G is the disruption of orderly transcription termination (the polyU tail at the 3’ end of *sgrS*). Such readthrough transcription has previously been ascribed to decrease the efficiency of SgrS binding to Hfq [28,29].

## Discussion

The construction of a stochastic kinetic model including gene replication, transcriptional regulation, and the role of the Hfq chaperone protein demonstrates the utility of combining single cell experiments with stochastic modeling. The SgrS Regulatory Network is a noisy system characterized by small numbers of sRNA and mRNA, as well as gene copy numbers that vary from cell-to-cell. This leads to the population level heterogeneity that can then be used to parameterize a kinetic model for analysis of the role of specific molecular actors, such as the chaperone Hfq, and the effects of point mutation on sRNA silencing of mRNA.

The average number of Hfq hexamers present in an *E. coli* cell has been reported to be on the order of 1400 to 10000 (2 *μ*M - 15 *μ*M) [25,36,40,45,47]. It is worth noting that an extensive microfluidic-based, single-cell proteomics study that analyzed over 4000 individual *E. coli* cells grown in similar media conditions as our study [40] found a mean Hfq level of 1500. Additional immunoprecipitation and sequencing studies (by RIL-Seq) have shown the number of various individual mRNAs and sRNAs being bound to Hfq to range from 10s to 1000 in *E. coli* [26]. Thus, our prediction (from fitting) that a pool of approximately 250 Hfq (0.5 *μ*M) are available to bind with SgrS sRNA at any time in the simulation of sugar shock regulation seems reasonable.

In addition, our approach allowed us to characterize the rate of Hfq-SgrS association compared to values reported for Hfq stabilization of other regulatory sRNAs. If the pseudo first order Hfq binding rate *k_bind_* reported in Table 1 is converted to a bulk second order rate we obtain a binding rate of 1.5 × 10^5^ *M*^−1^ *s*^−1^ which agrees reasonably well with the reported value [36] of approximately 10^6^ *M*^−1^ *s*^−1^ for long RNAs binding to Hfq [14,21] (compared to the value of to 10^8^ *M*^−1^ *s*^−1^ for short, unstructured RNAs binding to Hfq [17]). SgrS is a relatively long sRNA with a length of 227 nucleotides (sRNAs have been observed with 37-300 nt [44]), therefore the slow sRNA-Hfq binding process that we describe does seem likely. We suggest that this could be due to RNA restructuring of SgrS [1,8,24,38,39] by Hfq in order to promote binding with *ptsG* mRNA. It is thought that cellular sRNA and mRNA are present in large excess over Hfq [43], so nearly all cellular Hfq hexamers are thought to be bound to RNA. Since cellular mRNA in *E. coli* are thought to be on the order of approximately 2000-8000 copies [5] (much greater than the highest measured SgrS sRNA value of 200) the available Hfq pool size that we present is representative of the relative competitiveness (and time-dependent cycling) of SgrS for Hfq relative to the other particles that interact with the chaperone.

The study of mutant U224G shows the importance of Hfq stabilization in the SgrS regulatory network as a whole and seems to corroborate previous findings [31] that highlight the importance of the polyU tail for Hfq association with SgrS. The substantial decrease of the Hfq-SgrS binding rate and increase in the related unbinding rate relative to the *ptsG* mRNA annealing rates further down the network obtained from fitting confirms this point (Table 2). The changes in the SgrS-ptsG mRNA annealing rates *k_on_* and *k_off_* seem to support conclusions from the wild-type cells that Hfq-SgrS binding may result in some restructuring of the sRNA that makes this a slow process. This may explain the lower efficiency in *ptsG* mRNA association observed in mutant U224G, since Hfq cannot bind SgrS as effectively due to mutation at the polyU tail. Therefore, the predicted restructuring of SgrS by Hfq necessary to facilitate *ptsG* mRNA association is also hampered, resulting in slower and less stable mRNA binding (a decrease in *k_on_* and an increase in *k_off_*).

While this work is useful in describing the role of Hfq in the SgrS regulatory network and in capturing the stochastic nature of regulation over a population of replicating cells, it does not consider the various other SgrS mRNA targets that may be present in a living cell under certain growth conditions. In addition, other factors such as sRNA recycling (*i.e*. SgrS not being co-degraded with its target mRNA) [36, 38], which have been proposed for some sRNA and are now under study for SgrS, were not included, but can be incorporated into the model.

In conclusion, by incorporating gene replication, stabilization by the chaperone protein Hfq, and transcriptional gene regulation of *sgrS* we have developed a kinetic model capable of describing the cellular heterogeneity observed in the *E. coli* sugar shock response network. Stochastic simulation of the kinetic model allows us to take full advantage of the single-molecule fluorescence data that illustrates cell-to-cell variability in a collection of hundreds of cells. While the post-transcriptional regulation and silencing of *ptsG* mRNA by the sRNA is the critical feature, accounting for gene replication, transcriptional regulation, and stabilization gives a more robust picture of the regulatory network as a whole. In addition, complexifying the model highlights the importance of stabilization by Hfq and chaperone proteins in general in RNA silencing networks and allowed for a prediction of the rate of association of SgrS and Hfq (as a slow process, characterized by restructuring), the effective available Hfq pool size for the SgrS regulon under sugar stress conditions, as well as an analysis of an SgrS point mutation in one of the presumed Hfq binding modules (the polyU tail). The model presented in this work establishes a framework for models analyzing other sRNA mediated gene regulatory networks, and can be extended to spatially-resolved models describing SgrS target search kinetics.

## Supporting Information

**Supplemental File 1 - Appendix**

## Acknowledgments

The authors acknowledge Dr. Marie Ma for participation in helpful discussions regarding SgrS stability. This work was supported by grants from National Institutes of Health (NIGMS Grant R01 GM112659 and R35 GM122569) and through the National Science Foundation Physics Frontiers Center: “The Center for the Physics of Living Cells” (CPLC) (NSF PHY 1430124).

## Supplemental File 1 Appendix

### 1 Effects of Varying Available Hfq Pool Size

The available pool of Hfq utilized in the model represents the fraction of cellular Hfq hexamers bound to SgrS as opposed to other targets and thus the relative binding strength of SgrS compared to other RNAs stabilized by the chaperone. Previous work [26] has shown that the typical number of Hfq bound to a given sRNA varies widely across sRNA species. If an even smaller pool of cellular Hfq is assumed to be available for SgrS binding under sugar shock conditions the average behavior of SgrS experimentally observed can be more exactly captured (Figure 9). However, this comes at a loss of the population level noise observed in the measured RNA distributions because fewer SgrS can be stabilized and so it decays on a much faster timescale, resulting in a loss of cell-to-cell variation. When additional Hfq is added to the available pool such as the 800 available in the simulations shown in Figure 9 the opposite behavior can be seen. SgrS exhibits greater population level heterogeneity, but with a less robust fit to the average behavior that is experimentally observed. We propose that this creates more noise because SgrS is less likely to be present in its free form and decays more slowly when it is associated with *ptsG* mRNA and Hfq (*k_on_* is small relative to *k_ds_*) than it would when it is not stabilized by Hfq (Figure 4, **rxn 2.2** versus **rxn 4** followed by **rxn 5**).

**Figure 9.**
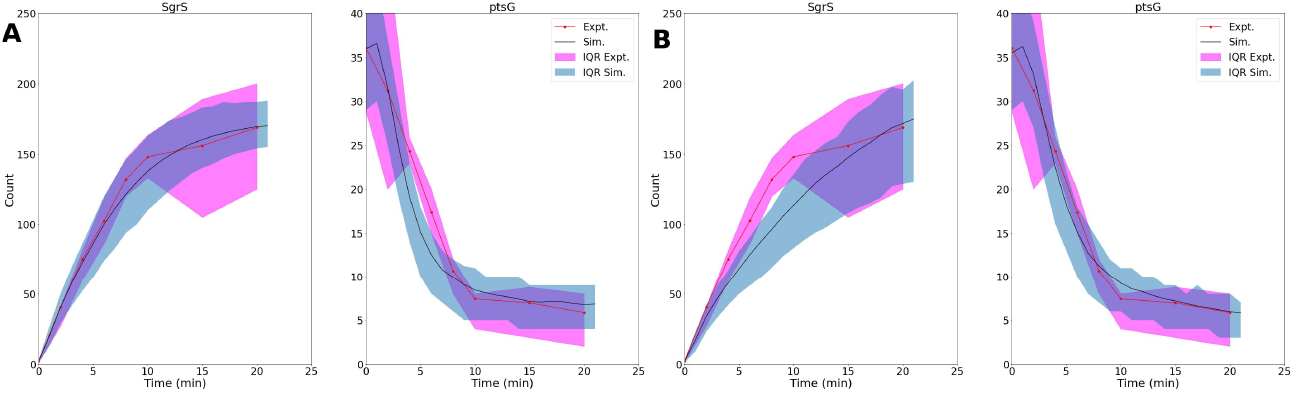
**(A)**: Trace and interquartile range (IQR) of SgrS sRNA and *ptsG* mRNA mRNA where simulations include a smaller pool of 200 Hfq available (versus 250 in main text simulations). While averages can be more tightly fit, the population level variation observed for SgrS is minimized even further from what is observed experimentally, including at long times post-induction. **(B)**: A similar plot of Trace and IQR with Hfq available pool size equal to 800. Here the population level variation is larger (especially at long times post induction), but the initial average traces are less well captured.

#### Effects of Initial Gene State

Of interest from a more technical standpoint, is the state of the *sgrS* genes at time = 0 minutes in the simulation. While, in principle these genes should be in the “OFF” state and unable to be transcribed since induction has yet to begin it is interesting to understand the effects of initial gene state on population level noise. Consider the following example, when all SgrS genes begin in the “ON” state. While the average behavior at times from 4 to 10 minutes is poorly captured, the RNA distributions are well-described at 15 and 20 minutes post-induction (Figure 10). This example assumes an immediate switch from the “OFF” to the “ON” state of the *sgrS* genes due to induction. While unrealistic when taken at face value, it is reasonable to assume the induction occurs on the order of seconds, since the amount of SgrS increases by a factor of 10 from its basal value by 2 minutes post-induction (Figure 5) and since binding of the SgrR activator for *sgrS* is mediated by binding to a small molecule (*i.e* sugar), which presumably takes some interval of time. The smaller *k_on,Ds_* and *k_off,Ds_* values (2.0 × 10 ^−3^ *s*^−1^ and 6.5 × 10^−4^ *s*^−1^ versus 3.0 × 10^−2^ *s*^−1^ and 9.5 × 10^−3^ *s*^−1^, respectively) used in Figure 4 Rxn 2.0 relative to those given in Table 1 then lead to a wider range of population distributions at late times due to longer dwell times (*i.e*. up to 5 minutes) for the *sgrS* gene in the “OFF” state compared to the a typical dwell time of less than 1 minute in the “OFF” state when the more appropriate regulatory values (3.0 × 10^−2^ s^−1^ and 9.5 × 10^−3^ s^−1^, based on the rapid increase in SgrS copy number from 0 to 2 minutes) are used for *k_on,Ds_* and *k_off,Ds_* respectively.

**Figure 10.**
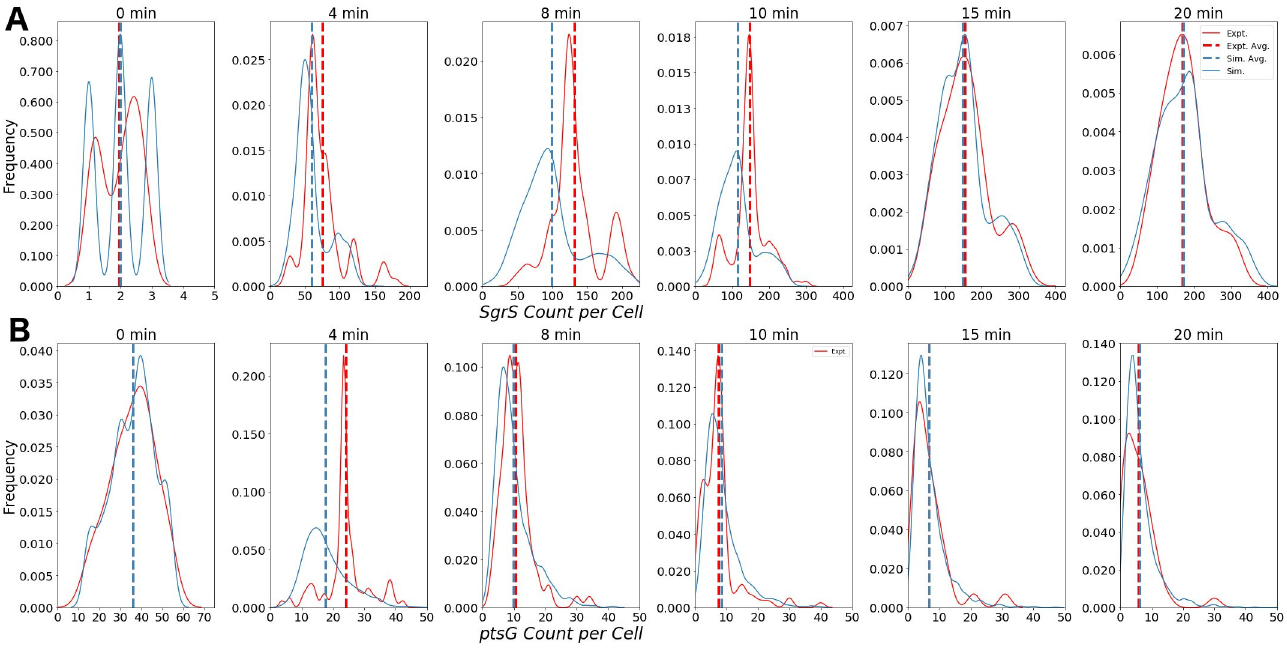
Distributions of **(A)** Wild-Type SgrS and **(B)** *ptsG* mRNA (bottom) at various time points from 0 to 20 minutes post-induction. Data from smFISH—STORM experiments (red, 100-200 cells per time point) and stochastic simulations (blue, 2000 cells per time point) are shown as kernel density estimates. Average copy number at each time point is displayed with dashed vertical lines.

#### Effects of Increased Cell Replicate Number

The number of *E. coli* cells that are simulated or have their RNA distributions experimentally measured is of great importance when considering a process characterized by stochasticity. A certain number of cells must be observed to accurately capture both the average behavior and cell-to-cell variability that emanates from a kinetic regulatory system [12,34,40].

Figure 11 shows the effect of number of cells measured on the average and standard deviation of the SgrS simulated at 20 minutes post-induction. The bootstrapping technique presented allows for the selection of an individual *E. coli* cellular replicate, with replacement, up to N cells. The vertical dashed line in each figure shows the expected average and standard deviation values produced from bootstrapping with N=85, the number of cells experimentally measured at time 20 minutes post-induction. This highlights the possible error in both mean copy number (5-10 copies) or population level variation (5-10 copies) that could be accrued due to insufficient sampling.

**Figure 11.**
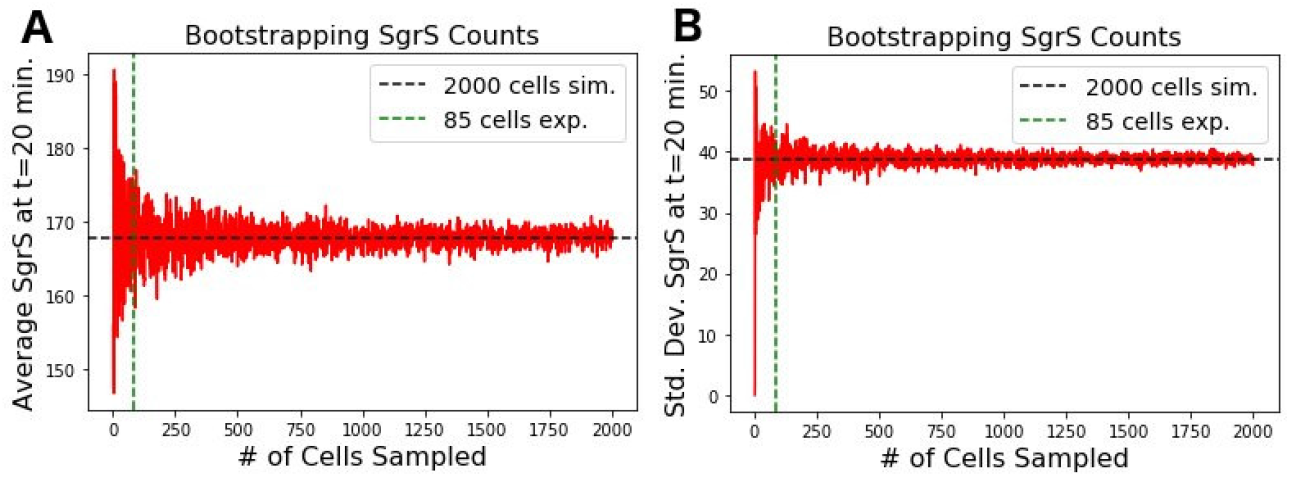
Bootstrapping of SgrS sRNA simulated at 20 minutes post sugar shock induction showing the variation in the **(A)** population mean and **(B)** population standard deviation with number of simulated cells sampled. The x axis gives the number of samples taken (N) with replacement out of a total 2000 independent simulation trajectories in the bootstrapping procedure. The vertical dashed line at N=85 shows the number of cells experimentally imaged at this time point. It takes several hundred to 1000 simulated cells before the SgrS mean and population level variation noise begin to relax to the calculated values.

#### Effects of SgrS Point Mutation On Regulatory Kinetics

In order to fit to mutant U224G the same parameters as for the wild-type cells were utilized other than the SgrS-*ptsG* mRNA binding and unbinding rates *k_bind_* and *k_unbind_* and the *ptsG* mRNA association rates *k_on_* and *k_off_*. The same gene state (high versus low gene copy number) percentages for *sgrS* and *ptsG* as for the wild-type cells were arrived at by fitting as well as the same “available” Hfq pool size of 250 hexamers. The distributions (as kernel density estimates) shown in Figure 12 for both SgrS and *ptsG* mRNA were obtained via the same fitting process described in the main text.

**Figure 12.**
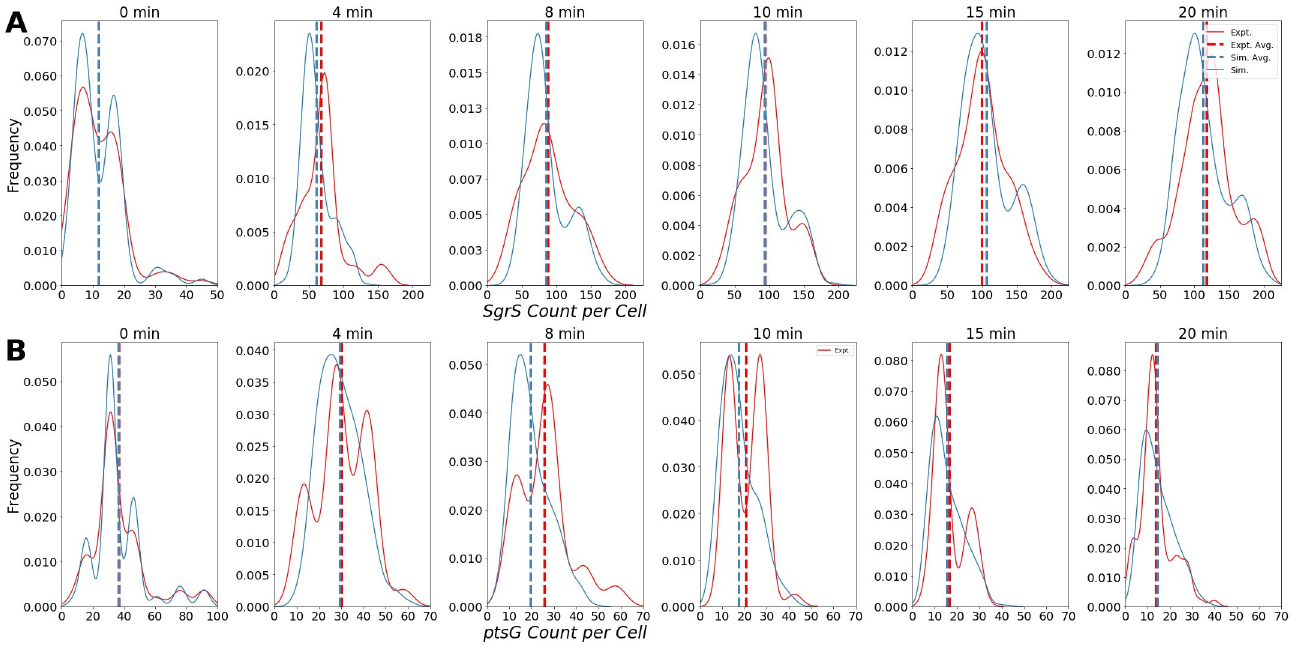
Distributions of the SgrS polyU tail mutant U224G for **(A)** SgrS and **(B)** *ptsG* mRNA at various time points from 0 to 20 minutes post-induction. Data from smFISH–STORM experiments (red, 100-200 cells per time point) and stochastic simulations (blue, 2000 cells per time point) are shown as kernel density estimates. Average copy number at each time point is are displayed with dashed vertical lines.

In order to focus on a point mutation that primarily showed a disruption in SgrS-Hfq association we sought a mutant in which SgrS secondary structure would not be significantly disrupted, leading to a higher free degradation rate of SgrS. In this way, we can isolate the effects of the point mutation on SgrS association to both the chaperone Hfq and its target *ptsG* mRNA individually. Via *in silico* folding using the RNA structure prediction tool mFold [49], we confirmed that the stability of the U224G with a ΔG of – 17.60 *kcal/mol* is unchanged from the predicted wild-type value of – 17.60 *kcal/mol*. The predicted U224G mutant structure also shows similar two stem loop structure (with loops of identical size) to that of the wild-type (Figure 13). Thus, an assumption that the measured wild-type ΔHfq degradation rate (see Main-text Section “Materials and Methods”) is appropriate for use as an SgrS-Hfq disassociation rate in fitting the U224G mutant data is reasonable.

**Figure 13.**
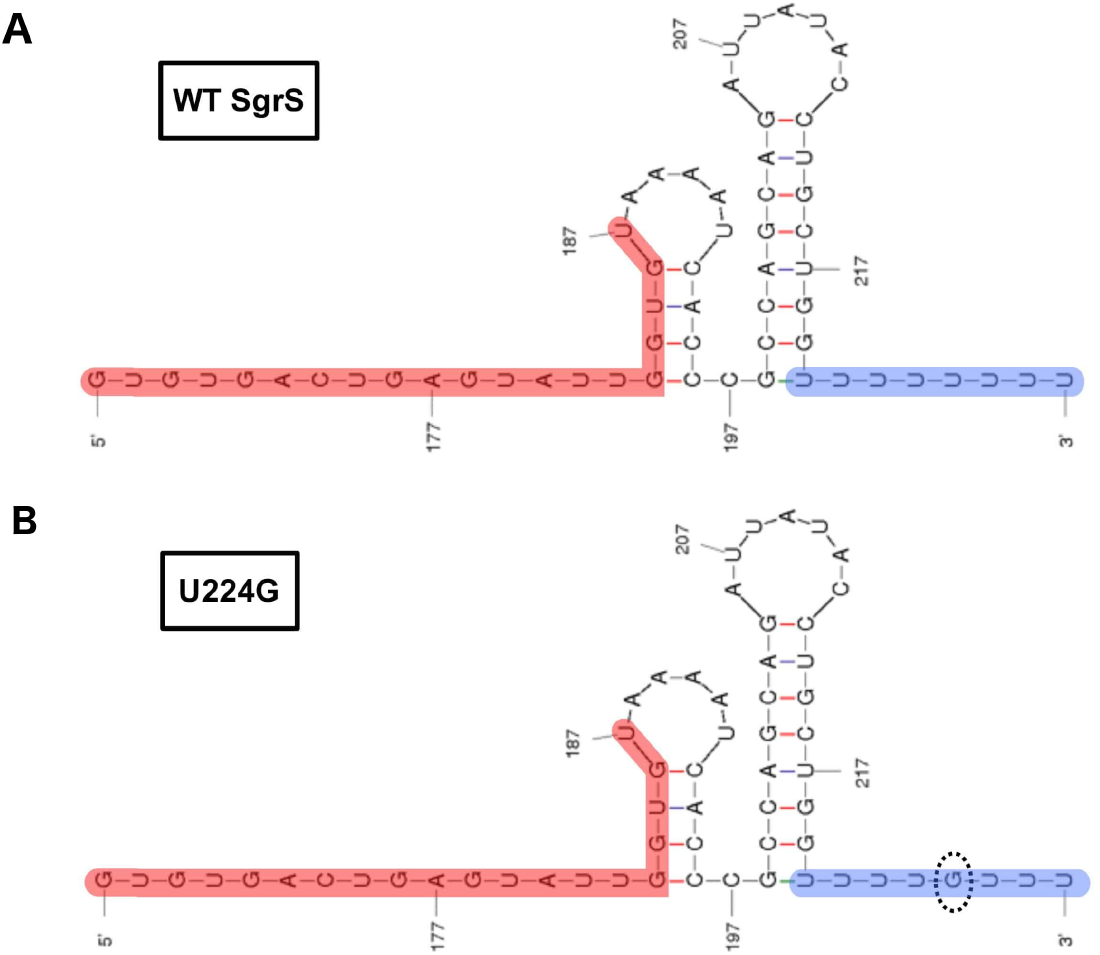
Flattened predicted sRNA structures for **(A)** wild-type (WT) SgrS as well as **(B)** the U224G mutant studied in this work obtained via mFold *in silico* folding. **Red**: the SgrS-*ptsG* mRNA baseparing region, **Blue**: the polyU tail, with the mutated residue circled in the U224G structure. The predicted structures show similar conformation as well as identical free energies (–17.60 *kcal/mol*), indicating that SgrS secondary structure is likely not significantly destabilized by the U224G point mutation in the polyU tail.

**Figure 14.**
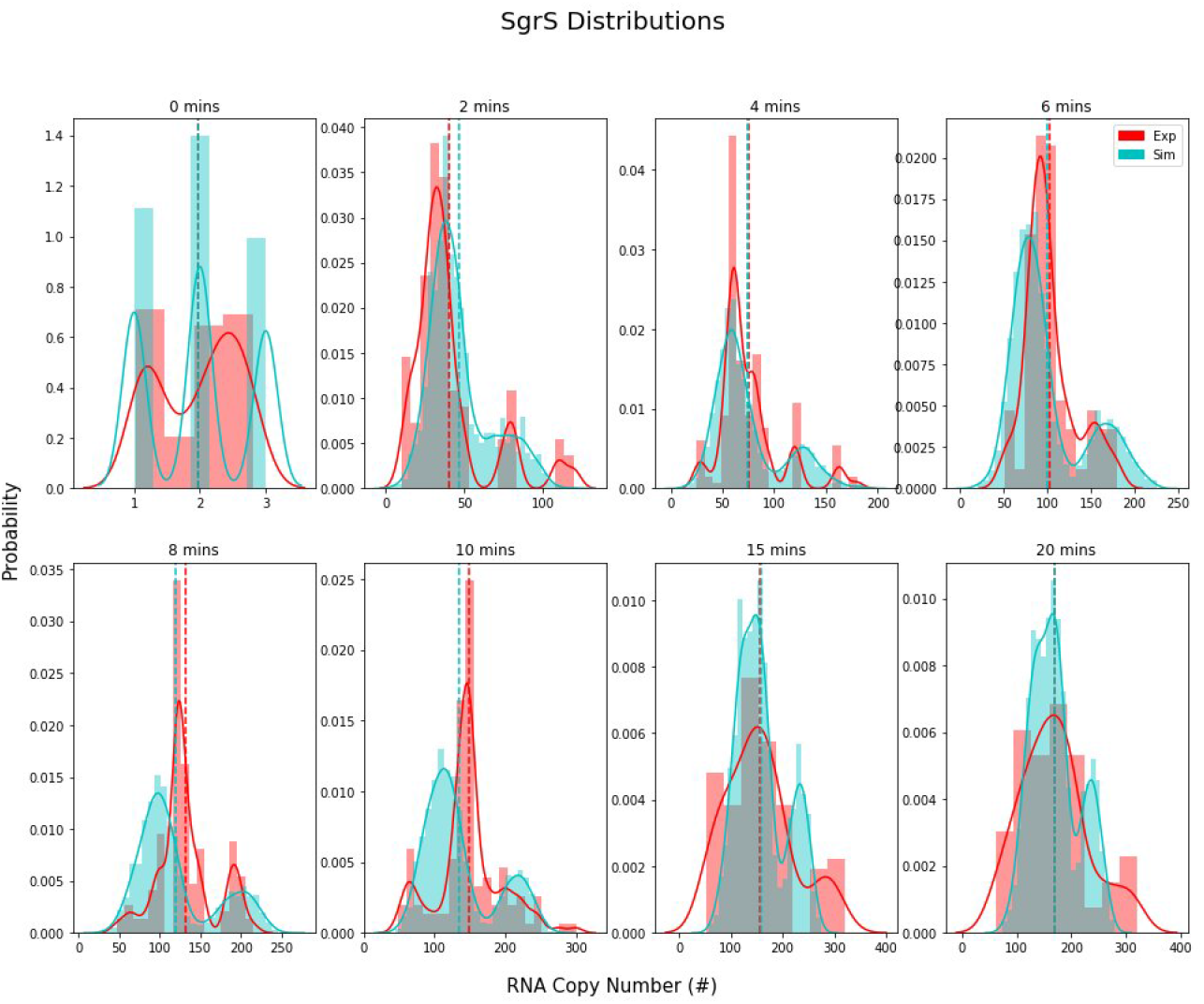
*ptsG* mRNA histograms, with experimental data in **red** and simulation data in **blue**. Scott’s normal reference rule was utilized to determine histogram bin widths.

**Figure 15.**
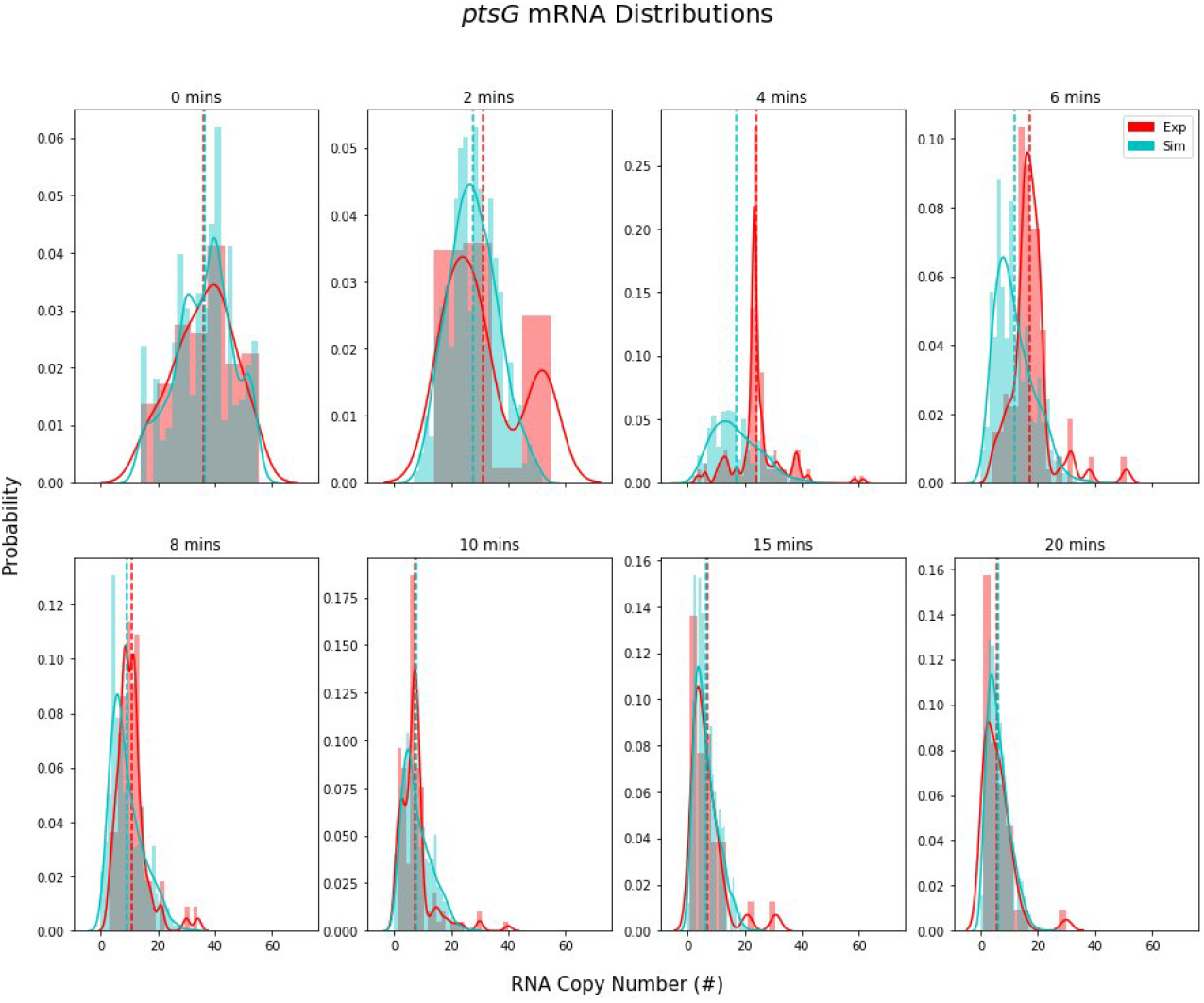
*ptsG* mRNA histograms, with experimental data in **red** and simulation data in **blue**. Scott’s normal reference rule was utilized to determine histogram bin widths.

#### Histograms of Experimentally Determined RNA Counts

The experimental data and simulated data shown in histogram form, prior to conversion to Kernel Density Estimates (KDEs) used in the main text and for fitting and analysis. Scott’s normal reference rule (Equation 3) was used to determine the bin width for the histograms of SgrS and *ptsG* mRNA at each time point. It is clear that the imposition of a kernel density on the experimental data, which was useful in constructing a fitting optimization scheme, has not fundamentally altered the character of the experimentally observed population level variation in SgrS or *ptsG* mRNA counts.

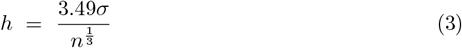

where *h* is the bin-width, *σ* is the sample standard deviation, and *n* is the number of samples (RNA copies).

